# State-dependent Online Reactivations for Different Learning Strategies in Foraging

**DOI:** 10.1101/2024.03.25.586512

**Authors:** Sangkyu Son, Maya Zhe Wang, Benjamin Y. Hayden, Seng Bum Michael Yoo

## Abstract

Reactivation of neural responses associated with navigation is thought to facilitate learning. We wondered whether reactivation is subject to contextual control, meaning that different types of learning promote different reactivation patterns. We trained macaques to forage in a first-person virtual maze and identified two distinct learning states prioritizing reward and information using unsupervised ethogramming based on low-level features. In orbitofrontal (OFC) and retrosplenial (RSC) cortices, representations of the goal, the path towards it, and recently traveled paths were strongly reactivated - online - during reward-prioritizing choices. During learning, reactivation of optimal paths increased in RSC after reward-prioritizing choices, and reactivation of uninformative paths decreased in RSC and OFC after information-prioritizing choices. Reactivation in OFC selectively covaried with ongoing RSC activity when prioritizing information; vice versa during prioritizing reward. These results highlight that cognitive states can drive learning and reactivation patterns can be tailored to the needs of the moment.

## INTRODUCTION

In our daily lives, we don’t simply attend to the world in front of us. Instead, our thoughts include recollecting the past and imagining the future. Far from idle daydreaming, these future and past-oriented thoughts are likely to play a crucial role in planning and learning. These mental events are recapitulated in neural processes as well, especially in the reactivation of neural activities. Most famously, recollection and imagination of travel are associated with the reactivation of place-sensitive firing in hippocampal place cells (Karlsson & Frank, 2009; Mou et al., 2022). This type of neural reactivation is directly linked to both learning and planning (Davidson et al., 2009; Dragoi & Tonegawa, 2010; Kuhl et al., 2013; Pfeiffer & Foster, 2013).

The most well-studied neural reactivation occurs during sleep (that is, offline) and predicts the degree of learning observed on waking (Diba & Buzsáki, 2007; M. A. Wilson & McNaughton, 1994). However, reactivation is also observed online, including between the trial intervals of behavioral tasks or while making a decision (Eldar et al., 2020; Jadhav et al., 2012; M. Z. Wang & Hayden, 2017). As with offline reactivation, online reactivation is thought to be associated with cognitive processes of retrospection, policy update, and planning, and appears to be associated with learning (Carr et al., 2011; Eldar et al., 2020; Liu et al., 2021).

Wakefulness is associated with striking variations in the cognitive state, which in turn have been associated with the prioritization of different objectives (Anderson & Perona, 2014; Pereira et al., 2020). Nonetheless, most accounts of neural reactivation treat it as having a single stereotyped form. We hypothesized, by contrast, that reactivation is subject to flexible moment-to-moment adaptive control, meaning that the decision-maker’s cognitive state can determine when and what form that reactivation takes. In particular, we hypothesize that different states will trigger different reactivation patterns tailored to the moment’s needs. Evidence for this hypothesis would not only show the influence of control over reactivation but would strengthen the evidence linking reactivation to specific learning processes.

To test this hypothesis, we focused on explore vs. exploit states. Natural decision makers in uncertain environments must trade-off between multiple objectives, namely, between efficient harvesting of reward (exploit states) and harvesting of information (explore states), which can be used to drive better information in the future (Cohen et al., 2007; Daw et al., 2006). Many studies have confirmed that the balancing of behavior between two states can occur in the highly constrained environment of the k-arm bandit task (Cohen et al., 2007; Ebitz et al., 2018; S. Wang et al., 2023; R. C. Wilson, Geana, et al., 2014). Some evidence suggests this behavior may be observed in more complex naturalistic situations as well (Marques et al., 2020). However, identifying it in highly complex foraging-like behavior remains a major challenge.

Here, we examined behavioral and population replay/preplay neural responses in two macaques performing a novel foraging task in a first-person virtual maze. Using only variables derived from elementary features of their behavioral repertories (and excluding choices and rewards), we identified two discrete latent states; these states have some face validity to explore and exploit states as they are generally defined. Simultaneously recorded population responses in the retrosplenial cortex (RSC) and orbitofrontal cortex (OFC) showed more neural reactivation patterns of the goal destination, the path to it, and recent past location trace when the subject focused on harvesting reward (that is, in the exploit-like state). In the explore-like state, where subjects chose the less visited paths, RSC and OFC showed decreased reactivation of the uninformative path. These replay/preplay patterns directly recapitulated the inferred learning strategies favored in each state. Specifically, the optimal path was reactivated more strongly in the exploit-like states, contrasting devaluation learning strategies for the explore-like state and reinforcement learning strategies for the exploit-like state.

## RESULTS

Two macaques separately performed a first-person perspective virtual reality (VR) foraging task based on a classic rat navigation task (Tolman & Honzik, 1930) (**Figure 1A** and **Supplemental Video 1**). They used a joystick to control their movement while we recorded neural activity from orbitofrontal (OFC) and retrosplenial (RSC) cortices (**Figure 1B**). The location of the jackpot reward (1 ml juice) was randomized daily; the starting position and initial facing direction were randomized on each trial. Each trial lasted until the subject reached the jackpot reward location.

**Figure 1.**
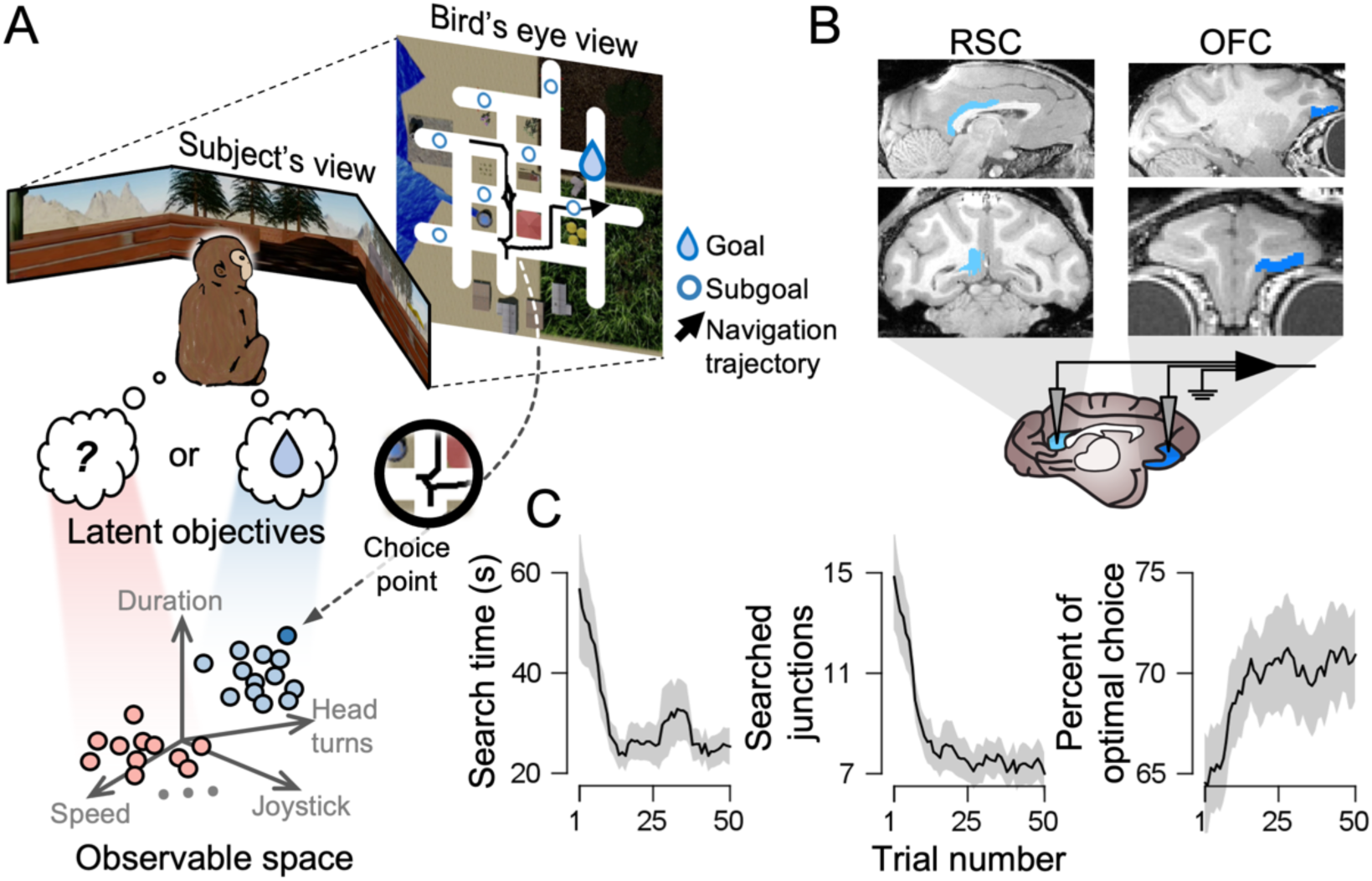
First-person virtual foraging task. **(A)** Top: subjects search for a jackpot reward at the goal location (with a tiny reward at subgoals) in virtual reality, teleported to a random location afterward. The goal location varied in each daily session and the starting position varied with each trial. Bottom: cartoon illustrating behavioral repertoires from observable low-level features at each choice point clustered according to latent objectives. **(B)** Recording site: retrosplenial cortex (RSC) and orbitofrontal cortex (OFC). **(C)** Improved performance over trials in time, distance, and optimality (reward-proximate). Shaded ribbons and crosshairs represent ±1 SEM.

Subjects completed each trial with a median duration of 12.3 seconds (subject P_(25%, 50%, 75%)_ = (10.0, 20.4, 38.0); subject S_(25%, 50%, 75%)_ = (5.7, 9.5, 18.7)). A key feature of our VR maze was its several choice points (junctions) within each trial. Each subject encountered a median of 5 choice points per trial (subject P_(25%, 50%, 75%)_ = (4, 6, 12); subject S_(25%, 50%, 75%)_ = (3, 4, 6)) and chose between 2-4 possible paths at each choice point. Importantly, these choices were not themselves rewarded, nor did they have a consistent spatial relationship with the reward. These choice points will be the focus of all subsequent analyses; we want to emphasize, however, that these choices are not trials. Instead, they are a sub-trial organizational structure imposed by the combination of the task and the subject’s own behavior. Subjects showed clear signs of learning across trials (**Figure 1C**). In particular, both search time and travel distance decreased with trial number in session (**Table 1**).

**Table 1.** Statistical values of linear trend analysis on foraging performance.

|  | Subject P |  | Subject S |  |
| --- | --- | --- | --- | --- |
| | $F_{(7,49)}$ | p | $F_{(8,49)}$ | p |
| Search time | 5.512 | 0.019 | 6.423 | 0.012 |
| Searched junctions | 8.456 | 0.004 | 35.672 | $5.420\text{e}^{-9}$ |
| % of optimal choice | 4.558 | 0.034 | 9.207 | 0.003 |

### Two latent learning strategies

We used unsupervised learning methods to see whether behavior at the choice point falls into distinct categories (**Figure 1A**, lower panel). Specifically, we chose twelve quantifiable low-level features of behavior: (i) head turn speed, (ii) head turn consistency, (iii) residence time at the choice point, speed (iv) at, (v) 500 ms before, and (vi) after the choice point, joystick press strength (vii) at, (viii) 500 ms before, and (ix) after the choice point, and acceleration (x) at, (xi) 500 ms before, and (xii) after the choice point. We excluded choice, choice outcome, and other task-related features to avoid using choice behavior to identify choice behavior (and thus avoid circular reasoning). Instead, we sought to use low-level variables to identify categorically different states (Berman et al., 2016; Voloh, Maisson, et al., 2023). We performed a t-distributed stochastic neighbor embedding (t-SNE) to visualize clusters (**Figure 2A** and **Supplementary** Figure 1). We found that behavior naturally falls into two, and not more, separate categories (**Figure 2B**; **Table 2**); this analysis is important for reducing the chance of reifying arbitrary clusters.

**Figure 2.**
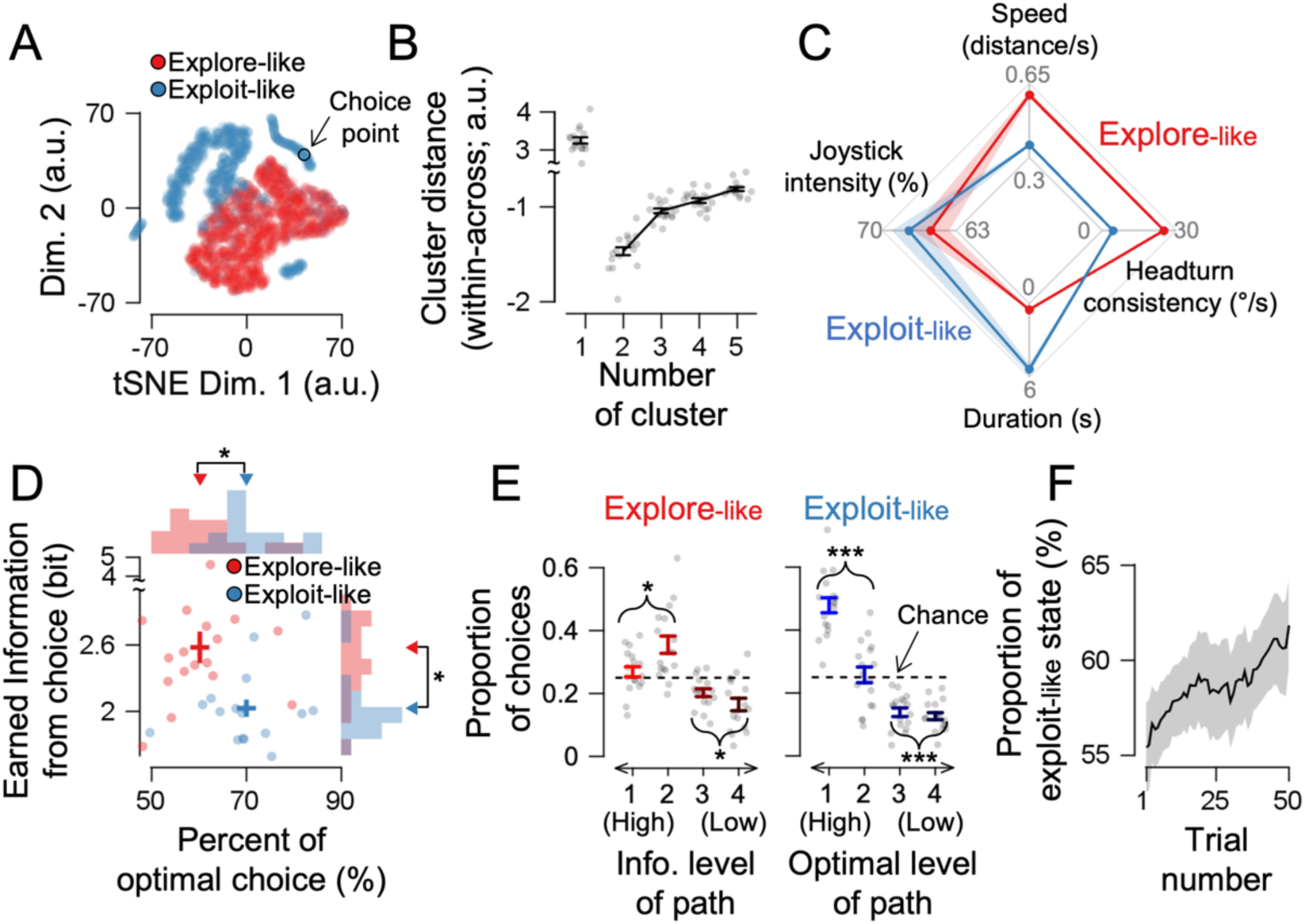
Two latent learning strategies. **(A)** Visualization of low-level features in t-SNE space (blue: exploit-like; red: explore-like). **(B)** K-means clustering performance was assessed for various cluster numbers (optimal K=2). **(C)** Four observable features of two states. **(D)** Decision outcomes: either optimal path in state 1 (named *exploit-like*) or informative (less explored) path in state 2 (named *explore-like*). **(E)** The proportion of path choices, ordered based on information in the explore-like state and optimality in the exploit-like state. **(F)** Over trials, the proportion of exploit-like states increased. Shaded ribbons and crosshairs represent ±1 SEM. Dots and stars indicate each session and statistical significance (*, p < 0.05; **, p < 0.01, *** p < 0.001).

**Table 2.**
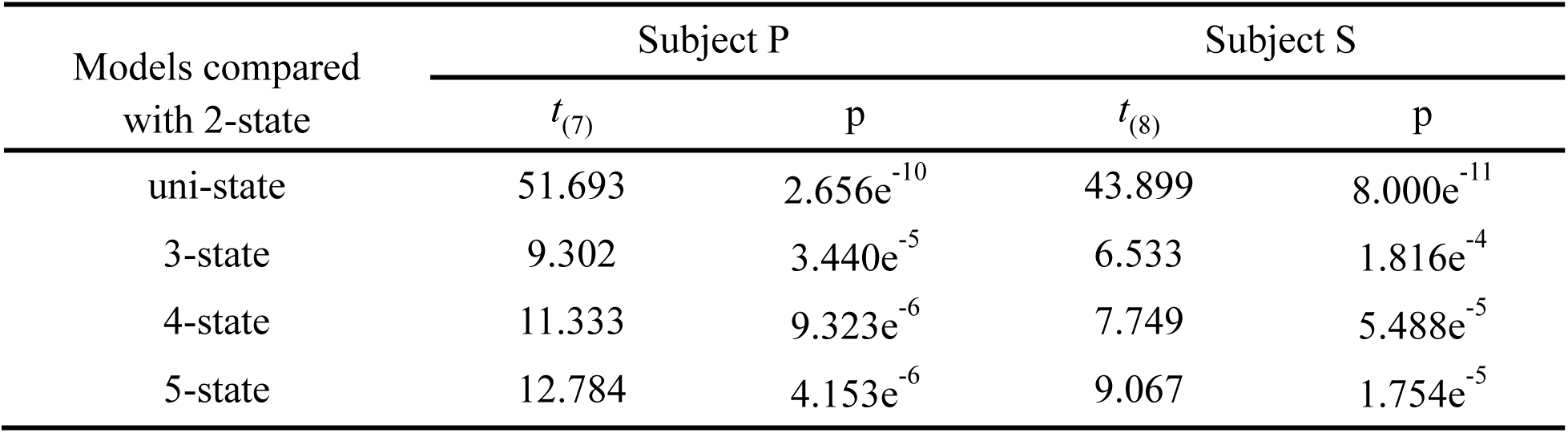
Statistical values of paired t-test on the performance comparison between the two-state k-means clustering model and N-state model.

In *state 1* (which we refer to as *exploit-like*), subjects moved slower and had more frequent head turns and stronger joystick presses after decision-making; these patterns were reversed in state 2 (which we call *explore-like*; **Figure 2C**; **Table 3**). We chose the term exploit-like because, in that state, subjects reliably chose the more optimal path, meaning it led more directly to the nearest jackpot reward. In the explore-like state, the subject reliably chose a more informative path, as indicated by selecting a less visited route (**Figure 2D**; subject P, *t*_info(7)_ = - 3.076, p_info_ = 0.017, *t*_opt(7)_ = 8.923, p_opt._ = 4.508e^-5^; subject S, *t*_info(8)_ = -3.439 p_info_ = 0.008, *t*_opt(8)_ = 3.164, p_opt._ = 0.013). Notably, performance in the explore-like state was still much better than in a random walk (**Supplementary** Figure 2; subject P, *t*_(7)_ = 14.578, p = 1.706e^-6^; subject S, *t*_(8)_ = 5.810, p = 4.002e^-4^). The path choices in the explore-like state were not random; among four path options, the subject consistently favored paths offering higher levels of information in the explore-like state (given that the information is an objective in the explore-like state) and chose less frequently for those with lower values (**Figure 2E** and **Table 4**). As subjects learned the maze, the proportion of exploit-like behavior increased within a session (**Figure 2F**; linear trend analysis; subject P, *F*_(7,49)_ = 13.263, p = 3.124e^-4^; subject S, *F*_(8,49)_ = 43.865, p = 1.162e^-10^), consistent with the idea of a shift from prioritizing information to reward, and in line with the conventional concept of explore and exploit, in which as the subject learned the maze, they derived less benefit from exploration.

**Table 3.**
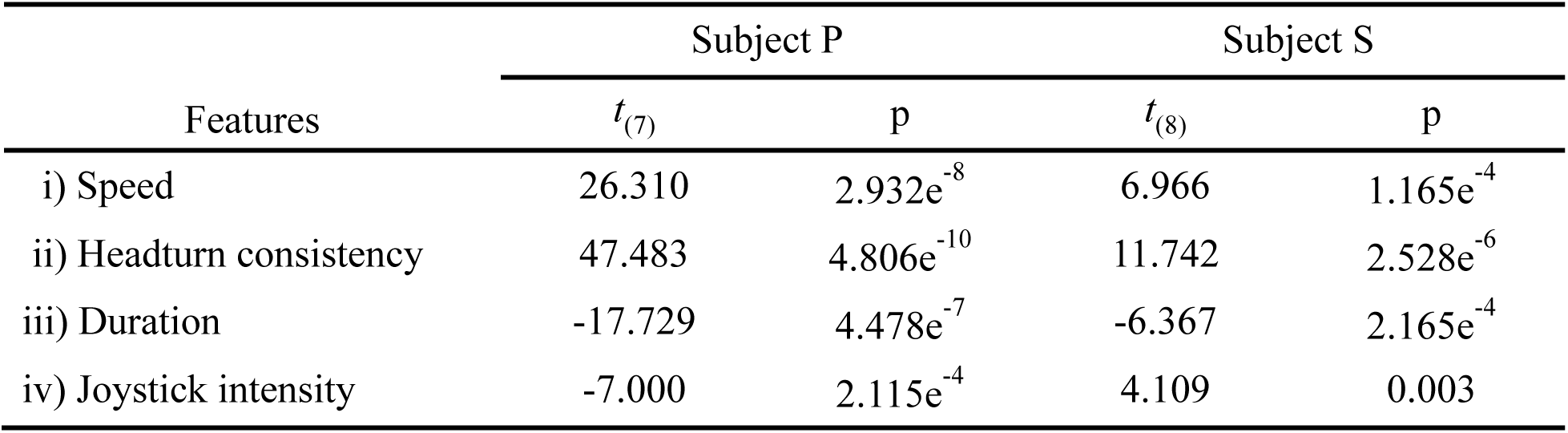
Statistical values of paired t-test on the low-level features of foraging behavior in the explore-like and exploit-like states.

**Table 4.** Statistical values of one-sample t-test on the proportion of chosen path.

| State | Level of<br>Info. / opt. | Subject P |  | Subject S |  |
| --- | --- | --- | --- | --- | --- |
| | | $t_{(7)}$ | p | $t_{(8)}$ | p |
| Explore-like | High | 4.696 | 0.002 | 2.569 | 0.033 |
|  | Low | -4.816 | 0.002 | -2.691 | 0.027 |
| Exploit-like | High | 14.417 | $1.839e^{-6}$ | 14.676 | $4.562e^{-7}$ |
| | Low | -17.257 | $5.389e^{-7}$ | -16.122 | $2.199e^{-7}$ |

Note, that the statistical test was done against the chance levels (=0.25).

### Neural reactivation in RSC and OFC

We recorded 655 (subject P) and 738 (subject S) neurons from the retrosplenial cortex (RSC) and 581 and 928 neurons (respectively) in the orbitofrontal cortex (OFC, **Figure 1B**). Although OFC is most well known for its role in reward processing (Padoa-Schioppa & Conen, 2017; Wallis, 2007), it also encodes the locations of future rewards during navigation (Basu et al., 2021; Wikenheiser et al., 2021) and the value of strategic exploration (Jahn et al., 2023). RSC is more traditionally associated with navigational computations, although its specific role remains unclear. Nonetheless, the preponderance of evidence suggests it may translate abstract spatial information into a navigational plan (Alexander et al., 2023; Miller et al., 2019; Vann et al., 2009). Neurons in RSC show reactivation during sleep (Feliciano-Ramos et al., 2023) and online (Staresina et al., 2013); it is not known whether neurons in OFC show reactivation during spatial navigation (although often the rewards were reactivated; Rushworth et al., 2011; Tsujimoto et al., 2009). We found a substantial proportion of neurons encoded goal position and spatial locations in both OFC (subject P, goal: 71.9% of neurons; location: 76.2%; subject S, goal: 66.6% of neurons; location: 60.2%) and RSC (subject P, goal: 70.2%; location: 73.8%; subject S, goal: 63.4%; location: 57.8%; **Supplementary** Figure 4).

We found clear evidence of reactivation in both OFC and RSC. Specifically, we found reactivation of the goal state, meaning that neurons showed activation patterns at the choice point that recapitulates the activation observed at the goal. To quantify goal state reactivation, we performed a vector alignment between normalized neural population activities at the moment of reaching the goal location and at the current junction with distinct states (**Figure 3A**). Reactivation was stronger in the exploit-like state than the explore-like state for both RSC and OFC (**Figure 3B**; **Table 5**). Note that in both states, we focused on cases where subjects chose the optimal path so the conditions were matched for reward expectations (see **Methods** for details).

**Figure 3.**
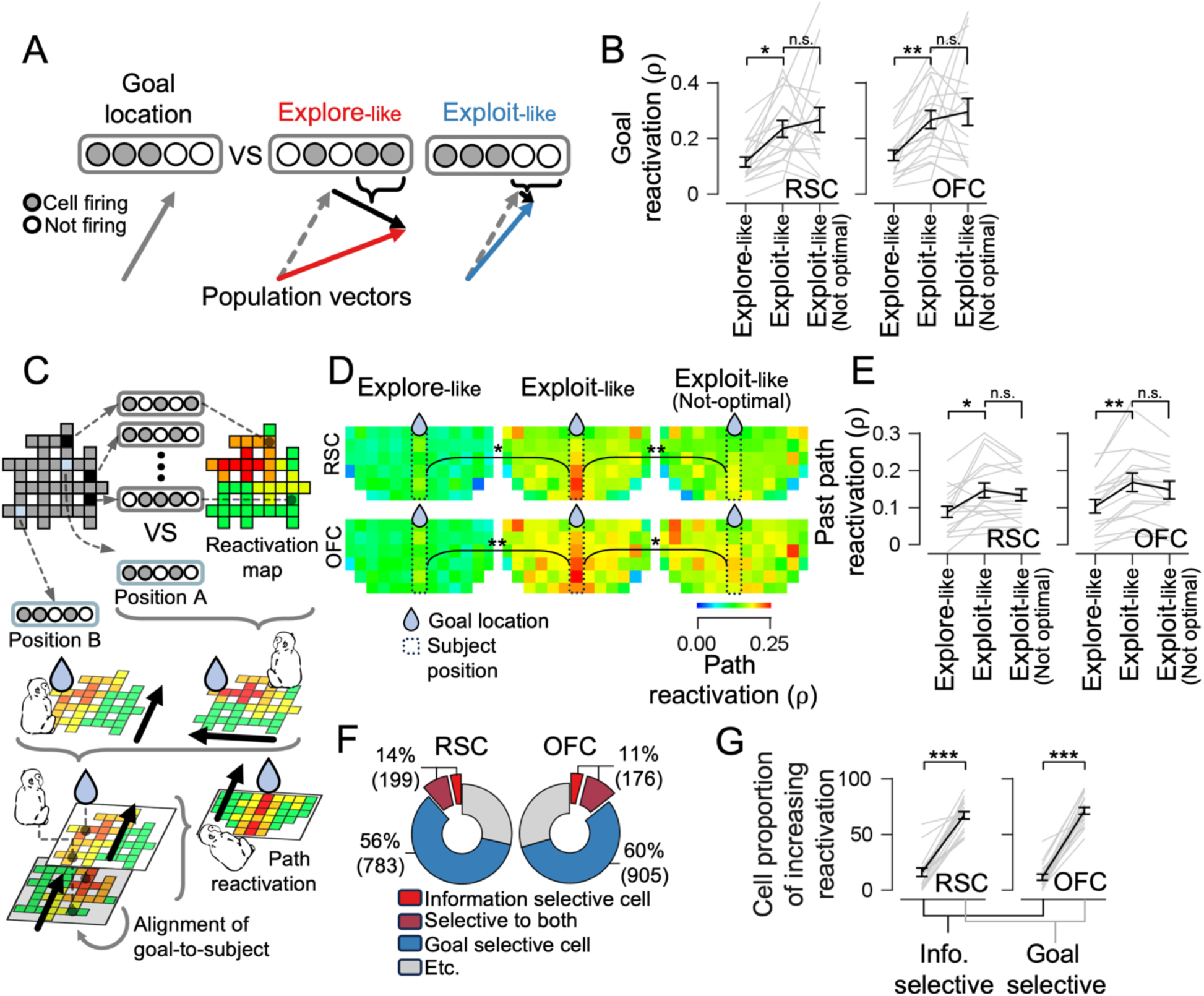
Neural reactivation of goal and path in RSC and OFC. **(A)** Goal reactivation. If the firing pattern of reaching the goal location is similar to that in the exploit-like state but not in the explore-like state, the population vector in the exploit-like state will be more aligned. **(B)** Reactivation is greatest in the exploit-like state. **(C)** Path reactivation. The firing pattern at one position was compared with that of all other possible junctions and corridors, organized into a maze layout. The reactivation maps were rotated to align to the various directions from different positions of the subject to the goal location. **(D)** In the exploit-like state, reactivation was higher in the path from the subject’s position to the goal location (dotted squares), except in cases where the subsequent choice was not optimal. **(E)** In the exploit-like state, the past path (the corridor and the junction just passed) showed a higher level of reactivation. **(F)** Of the recorded cells, 11 to 14% of the cells showed selective firing rate changes for information. **(G)** The cell proportion showed increased reactivation after being included in the reactivation analysis. Gray lines represent each session. The error bar represents ±1 SEM. Stars indicate statistical significance (*, p < 0.05; **, p < 0.01; ***, p < 0.001; n.s., not significant).

**Table 5.**
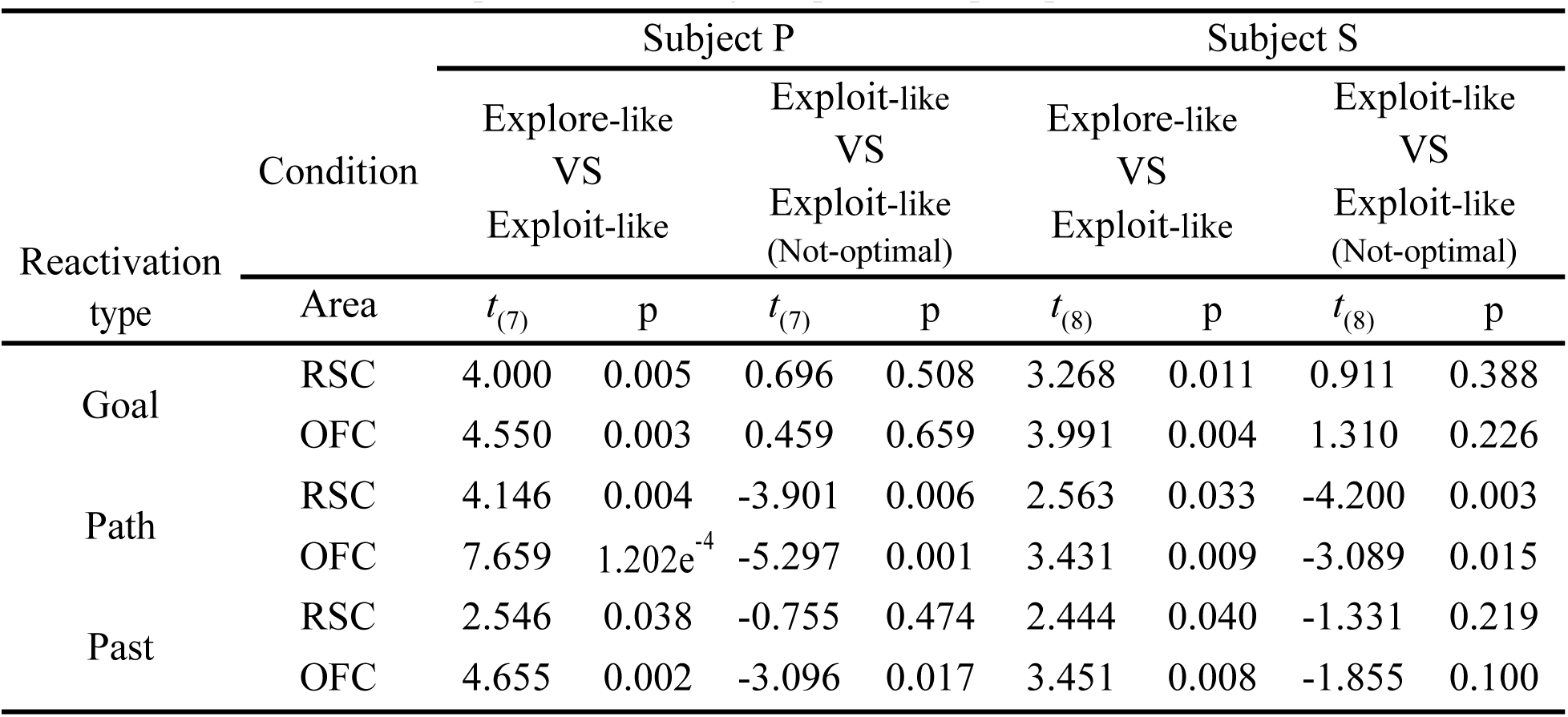
Statistical values of paired t-test on goal, path, and past path reactivation.

We also found reactivation of the path between the subject’s current location and the final destination (*path reactivation*; Brown et al., 2016; Howard et al., 2014) (**Figure 3C**). The reactivation map was measured by comparing the firing pattern at the current and all the other locations and then rotated to make them consistent in direction toward the goal locations (see **Methods**). Stronger path reactivation was observed in the exploit-like state than in the explore-like state for both RSC and OFC (**Figure 3D**; **Table 5**). We also found the stronger reactivation of recently occupied locations in the exploit-like state (i.e., *past reactivation*; Davidson et al., 2009) (**Figure 3E**; **Table 5**). One interpretation of these results is that exploit-like states more effectively bridge past experience and reward-related future plans (Byrne et al., 2007; Hahamy et al., 2023; van der Meer et al., 2010).

We next examined whether the degree of reactivation predicts subjects’ choice behavior. Surprisingly, we found that suboptimal (less reward-directed) and optimal choices were associated with equally large goal and past reactivation (**Figure 3B and E**; **Table 5**). However, the reactivation of the path to the reward was significantly reduced in suboptimal exploit-like choices in both regions (**Figure 3D**; **Table 5**). This finding is consistent with the hypothesis that suboptimal choices may arise from reactivating the goal without proper reactivation of the path to the goal (Howard et al., 2014).

We subsequently asked whether the vector misalignment was attributable to the properties of individual neurons (Genzel et al., 2020). Given that the population vector in the explore-like state had a larger misalignment with the goal state, we hypothesized that the neurons selective to informativeness were responsible for reduced goal reactivation. We first found that 11-14% of neurons had an activity that was significantly correlated with information gain at each choice point (**Figure 3F**). Notably, the majority of the neurons that were selective to informativeness also exhibited selectivity to the goal location. A smaller number of information-selective cells, compared to the goal-location selective cells, showed an increase in goal reactivation when the cell was included in the reactivation analysis (**Figure 3G**; permutation; subject P, t_RSC(7)_ = 5.794, p_RSC_ = 6.773e^-4^, t_OFC(7)_ = 8.481, p_OFC_ = 6.265e^-5^; subject S, t_RSC(8)_ = 5.220, p_RSC_ = 8.018e^-4^, t_OFC(8)_ = 5.201, p_OFC_ = 8.209e^-4^). This suggests that the neural activity may diverge away from encoding reward-related reactivation due to individual neuron’s selectivity to informativeness.

### State-selective navigational learning by distinct reactivation

We hypothesized that the explore-like and exploit-like states might be associated with two distinct learning patterns (**Figure 4A**). Specifically, we reasoned that, in the explore-like state, subjects might show devaluation-based learning (Daw et al., 2005; R. C. Wilson, Takahashi, et al., 2014; Wrase et al., 2007) (i.e., rule out uninformative options); in other words, they might learn more from an uninformative choice than an informative choice. In this case, in the subsequent visit, they should opt for a path more aligned with the information-prioritizing objective. Indeed, we found just that: subjects selected a more informative path after an uninformative choice in the previous visit (**Figure 4B**, left; subject P, *t*_(7)_ = -6.784, p = 2.567e^-4^; subject S, *t*_(8)_ = -3.751, p = 0.005). This suggests that subjects were more sensitive to uninformative paths than informative ones. At the same time, subjects were not sensitive to their choice being optimal or not in the explore-like state, since subjects chose paths with a similar frequency of visits, irrespective of whether the previous choice in the explore-like state led to a closer (optimal) or farther location (suboptimal) (**Supplementary** Figure 5, left; subject P, *t*_(7)_ =-1.555, p = 0.163; subject S, *t*_(8)_ = -1.664, p = 0.134). RSC and OFC activity recapitulated devaluation-based learning patterns: they showed reduced neural reactivation of the previously uninformative path while leaving the informative one unaffected (**Figure 4C**; **Table 6**).

**Figure 4.**
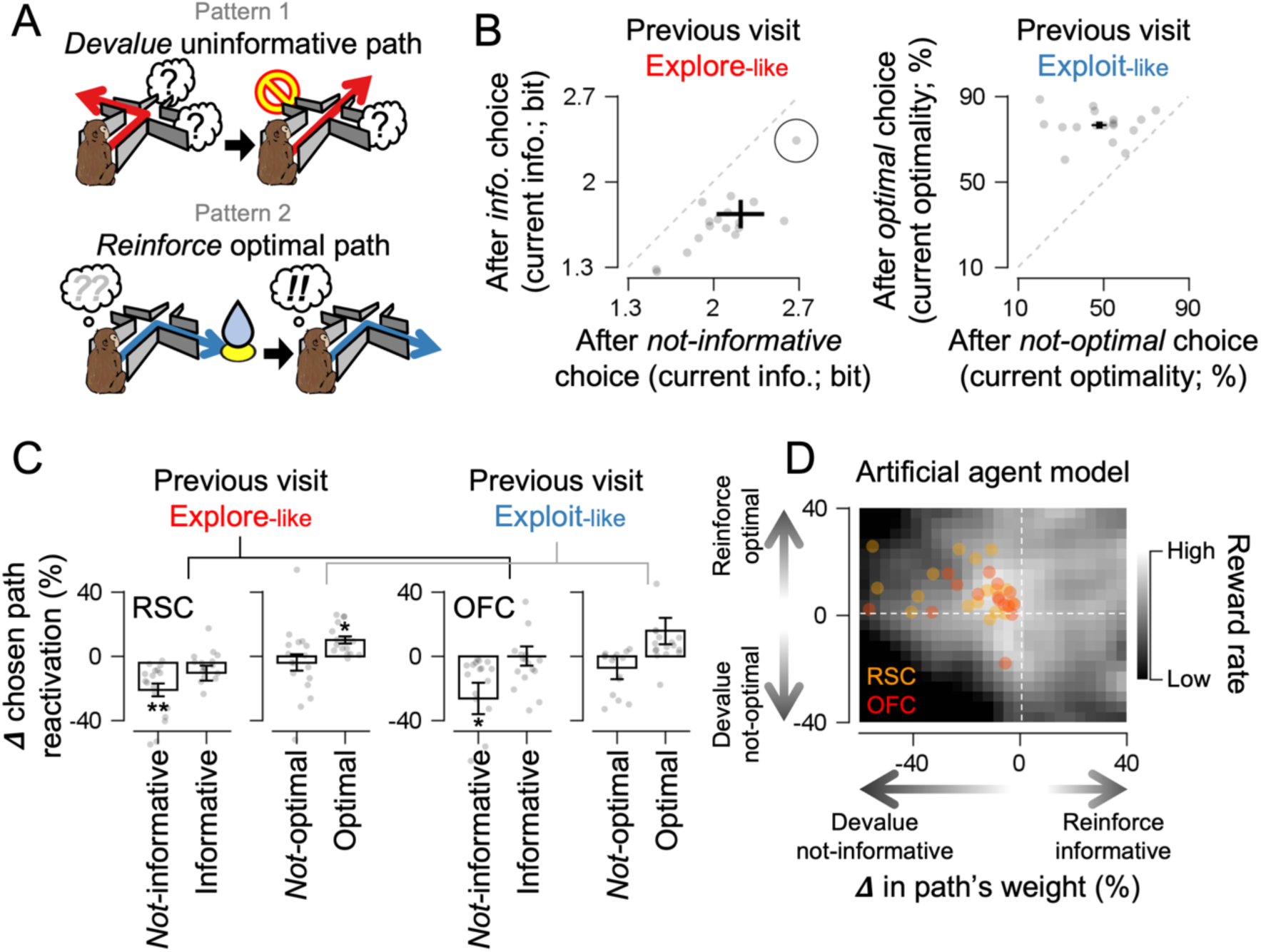
State-selective navigational learning by distinct reactivation. **(A)** If the choice was uninformative, the specific path might be devalued in the explore-like state. If the choice was optimal, the path might be reinforced in the exploit-like state. **(B)** Upon uninformative choice in an explore-like state, in the subsequent visit, the subject avoided the same path and gained information more (left). In contrast, upon optimal choice in an exploit-like state, in the subsequent visit, the subject chose the same path and gained higher optimality (i.e. reward-approaching) (right). The black circle denotes an outlier for visualization purposes. **(C)** In RSC and OFC, the corresponding path reactivation decreased following an uninformative exploration in the previous visit, whereas it increased in RSC after an optimal decision in the previous exploitation. **(D)** The reward rate of artificial agents foraging in the same maze. The agent’s choice was proportional to weight, which was updated (devaluing or reinforcing) based on informativeness and optimality. Percent changes in the reactivation of uninformative paths and optimal paths in the RSC and OFC (from (C)) were superimposed. Other symbols are identical to the former figures.

**Table 6.** Statistical values of one-sample t-test on the percent change of the chosen path reactivation after a certain condition.

| Previous<br>state | Previous<br>choice | Subject P |  |  |  | Subject S |  |  |  |
| --- | --- | --- | --- | --- | --- | --- | --- | --- | --- |
|  |  | RSC |  | OFC |  | RSC |  | OFC |  |
| | | $t_{(7)}$ | p | $t_{(7)}$ | p | $t_{(8)}$ | p | $t_{(8)}$ | p |
| After<br>Explore-like | *Not-*<br>informative | -4.131 | 0.004 | -3.618 | 0.008 | -4.176 | 0.003 | -2.328 | 0.048 |
|  | Informative | -2.979 | 0.021 | -2.417 | 0.046 | -1.630 | 0.141 | 0.760 | 0.469 |
| After<br>Exploit-like | *Not-*<br>optimal | 1.041 | 0.332 | -0.375 | 0.718 | -1.022 | 0.336 | -0.977 | 0.357 |
|  | Optimal | 3.243 | 0.014 | 5.063 | 0.002 | 3.072 | 0.015 | 1.360 | 0.210 |

While the explore-like state was associated with devaluation-based learning, the exploit-like state was associated with standard reinforcement learning (reinforcement-based learning). Specifically, when a path that had been previously chosen during the exploit-like state led to a closer location to the reward (i.e., it was a more optimal choice), subjects were more likely on subsequent visits to choose the optimal path again (**Figure 4B**, right; subject P, *t*_(7)_ = 4.470, p = 0.003; subject S, *t*_(8)_ = 4.835, p = 0.001). Consistent with this strategy, reactivation of the previously optimal path was enhanced in RSC, but reactivation of the suboptimal path was unchanged (**Figure 4C**; **Table 6**). Thus, explore-like and exploit-like states were associated with distinct learning strategies, which corresponded to distinct reactivation patterns.

We further hypothesized that the combination of two distinct learning patterns served the purpose of maximizing reward intake. To test this, we used an agent-based modeling procedure to simulate foraging in a virtual environment (similar to the approach used in Banino et al., 2018). We measured the reward rate (ml per traveled distance) of an artificial agent in the same foraging task (**Figure 4D**). We found that the agent’s path choice was proportional to weight, where the weight of the chosen path was updated using various combinations of devaluing or reinforcing patterns for the subsequent visit. The agent was most rewarded when simultaneously 8% devaluing the uninformative path and 12% reinforcing the optimal path. These proportions matched with the values of two learning patterns shown in RSC and OFC (RSC: median devaluation at 15.7%, reinforcement at 9.2%; OFC: median devaluation at 8.3%, reinforcement at 6.6%). This simulation result indicates that the two learning patterns we observe in the ratio are consistent with strategic reward optimization, given the constraints imposed by uncertainty.

### Cross-areal interaction is state-dependent

As the hippocampal-OFC interaction (Wikenheiser & Schoenbaum, 2016), including RSC-OFC (Alexander et al., 2023), is known for connecting reward and space, we found that when reactivation was higher in OFC, it was also higher in RSC, and vice versa (**Figure 5A**, mean Spearman correlation coefficient = 0.478; subject P, t_(7)_ = 8.477, p = 6.282e^-5^; subject S, t_(8)_ = 7.959, p = 4.527e^-5^). Based on this finding, we hypothesized that two areas might interact in a state-specific manner. To test this hypothesis, we calculated multiple linear regression between the individual cell firing rate of one area and the other area’s goal reactivation separately for each state; the collective regression coefficients were represented as vectors (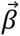_explore-like_ and 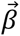_exploit-like_; **Figure 5B**). In the explore-like state, we found that RSC neurons’ firing was associated with OFC ensemble reactivation, while OFC activity was not associated with RSC’s reactivation (**Figure 5C**; **Table 7**). In the exploit-like state, the reverse relationship was true – OFC neurons’ activity was associated with RSC ensemble reactivation, but RSC was not associated with OFC’s reactivation (**Figure 5C**; **Table 7**).

**Figure 5.**
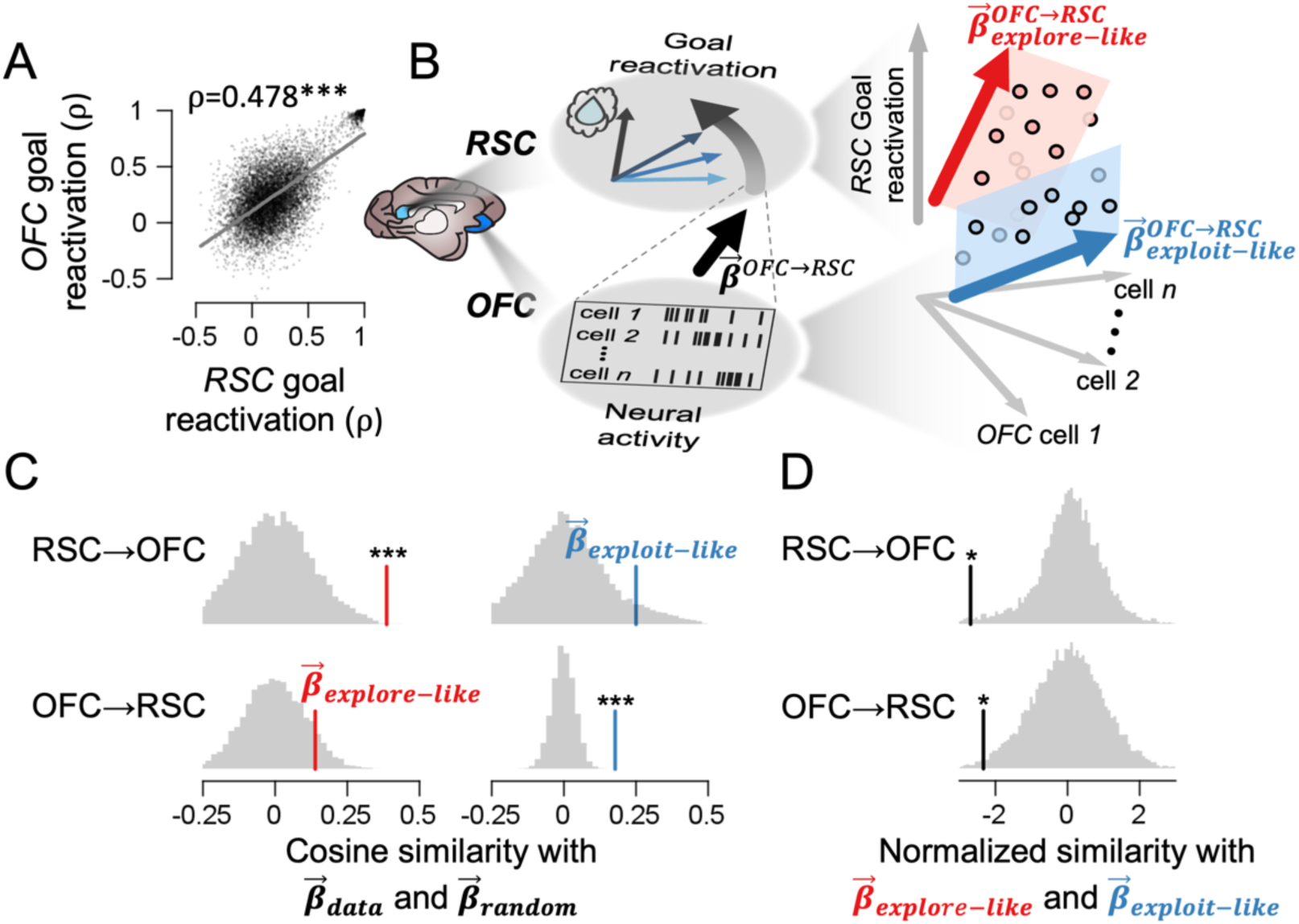
Cross-areal interaction is state-dependent. **(A)** The goal reactivations of the two regions were correlated with each other. The dots represent choice points. **(B)** To unveil the relationships between the firing rate of one region and the change in the other region’s goal reactivation, multiple linear regression coefficients were measured (represented as vectors; 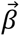*_explore-like_* and 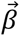*_exploit-like_*). **(C)** 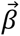*_explore-like_* exhibited greater similarity with resampled vectors (red vertical lines; showing consistency of 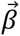) than random vectors (background gray distributions), specifically when the RSC firing rate was projected onto OFC reactivation, not vice versa (left panels). In contrast, 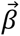*_exploit-like_* exhibited greater similarity with resampled vectors (blue vertical lines) than random vectors (background gray distributions), exclusively when OFC firing rate was projected onto RSC reactivation, not vice versa (right panels). This indicates two regions interacted state-selectively. **(D)** 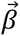*_explore-like_* and 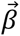*_exploit-like_* were less effective on each other (vertical lines) compared to randomly selected two vectors (background gray distribution) during across-region interaction. Analysis was done after all cells from two subjects were combined. Stars indicate statistical significance (*, p < 0.05; ***, p < 0.001).

**Table 7.** The p-values of the permutation test on the regression coefficient vector between one area’s firing rate and the other area’s goal reactivation.

|  | Combined |  | Subject P |  | Subject S |  |
| --- | --- | --- | --- | --- | --- | --- |
|  | RSC | OFC | RSC | OFC | RSC | OFC |
|  | → | → | → | → | → | → |
|  | OFC | RSC | OFC | RSC | OFC | RSC |
| Explore-like VS random | 1.562e<sup>-4</sup> | 0.092 | 0.065 | 2.439e<sup>-6</sup> | 9.354e<sup>-7</sup> | 0.110 |
| Exploit-like VS random | 0.065 | 7.047e<sup>-7</sup> | 0.231 | 1.409e<sup>-7</sup> | 0.006 | 0.002 |
| Explore-like VS Exploit-like | 0.021 | 0.012 | 0.011 | 1.960e<sup>-5</sup> | 0.029 | 0.030 |

Note, (area A)→(area B) indicates the regression coefficient vector was estimated using area A's firing rate to predict area B's goal reactivation. When one subject showed significance while the other did not, the statistical tests were performed after all cell data were collapsed over two subjects (combined).

Next, we hypothesized whether the inter-areal association in one state is effective in the other state’s association. That is, the two reactivation vectors for each state (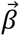_explore-like_ and 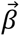_explore-like_) in a single brain area might correlate with each other. To test this hypothesis, we projected the one regression coefficient vector onto its conjugate (i.e., cosine similarity between 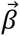_explore-like_ and 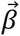_exploit-like_ vectors). The association of RSC neurons’ firing with OFC’s reactivation in the explore-like state was independent of that in the exploit-like state (i.e., null projection of one vector onto the other), as was the association of OFC neuron’s firing with RSC reactivation (**Figure 5D**; **Table 7**). This pattern indicates that one area interacting with the reactivation of another area in a particular state was both independent and selective. Moreover, these results indicate that, while there is bidirectional communication between RSC and OFC, the content of the information is not necessarily reciprocal.

## DISCUSSION

We investigated neural signatures of replay and replay in populations of neurons in OFC and RSC while macaques performed a virtual foraging task. We found signatures of reactivation of relevant paths and destinations in both regions, and, critically, that the form of reactivation varies according to its presumed functional role. Specifically, we used a novel bottom-up ethogramming process to identify two latent states associated with different learning patterns one in which subjects preferred visiting less explored paths (explore-like), while the other involved a preference for reward rate-maximizing path (exploit-like). Reactivation patterns in both regions differed with state, and, indeed, showed the pattern expected if they were tailored to the learning demands of the state. Specifically, in the exploit-like state, we observed a reactivation pattern that recapitulates the goal, the path to the goal, and the recent path. In the explore-like state, reactivation was weakened for the frequently-visited, thus uninformative, path, whereas, in the exploit-like state, the reactivation reinforced the path towards the goal in RSC. These results support the hypothesis that reactivation is regulated in a way that is tailored to the needs of the state.

Learners often optimize other objectives beyond mere exploitation (i.e., maximization of local rewards), including curiosity (i.e., maximization of information; Kidd & Hayden, 2015; Poli et al., 2024) and empowerment (i.e., maximization of the number of future outcomes; Brändle et al., 2023; Klyubin et al., 2005). Our results demonstrate that distinct objectives in each learning process involve distinct types of neural reactivations, suggesting that neural reactivation could serve as a generalizable neural substrate for various learning objectives.

Our results highlight the role of OFC in navigational planning. It is most often assumed that navigation is the domain of a specialized and localized circuit that consists of the hippocampus and adjacent areas, such as the RSC. Indeed, the important work that has established the validity of preplay and replay is generally localized to these regions (Byrne et al., 2007; Miller et al., 2019; van der Meer et al., 2010; Vann et al., 2009). However, some recent studies suggest that the OFC, which has strong bidirectional connections with hippocampal regions, may also encode spatial information related to goals, suggesting it contributes to navigation as well (Basu et al., 2021; Maisson et al., 2023; Wikenheiser et al., 2021). Our results confirm these findings and extend them by showing that not only is the present position encoded, but also OFC shows reactivation. These results, in turn, challenge the narrow view that focuses on the role of OFC solely in value representation and comparison, and suggests it participates in more complex and especially in grounded processes.

These results also confirmed the predicted role of RSC. Due to its anatomical adjacency to the hippocampus, most theories on RSC’s function focused on its putative role in navigation (Alexander et al., 2023; Vann et al., 2009), including in linking the past trace to future planning (Byrne et al., 2007). Although studies have supported this idea using rodents (Miller et al., 2019; van der Meer et al., 2010), uncertainty has still remained due to scarce recordings in primates. Our results validate that the RSC in primates is involved not only in replay/preplay for navigational planning (as with OFC) but also in reallocating the reactivation strength across option paths with learning context. Together, these results strengthen the empirical support for RSC’s contribution to learning in spatial navigation.

Modern computation and data-heavy approaches to ethogramming have generally confirmed the classical ethological idea that behavior is naturally divided into discrete behaviorally relevant states (Berman et al., 2016; Pereira et al., 2019; Wiltschko et al., 2015). This is also true in monkeys (Manea et al., 2024; Voloh, Eisenreich, et al., 2023; Voloh, Maisson, et al., 2023). These states are presumably associated with distinct cognitive repertoires that support the needs of that state. The distinction between explore and exploit states, which differentially emphasize information- and reward-seeking, is one of the most well-studied state-based divisions of cognitive behavior (Chen et al., 2021; Daw et al., 2006; Ebitz et al., 2018), although it has seldom been detected in naturalistic behavior (Marques et al., 2020). The differential role of information in these two states suggests that they may be associated with different types of learning; here, we confirm this idea in our own data and show that these forms of learning are associated with distinct reactivation patterns. Together, these findings illustrate that the brain regulates its learning strategy, and the neural processes support its particular needs of the state it is in. More generally, these results suggest that drawing broad assumptions from simple laboratory tasks may be limited to the state in which they were recorded and may not readily generalize to other ones. They, therefore, point us to the complex richness of naturalistic cognition (Yoo et al., 2021, 2020).

## Acknowledgment

This research was supported by IBS-R015-D1 (SS and SBMY), RS-2023-00211018 (SS and SBMY), MH129439 (BYH), and R01 MH125377 (BYH). The authors declare no competing financial interests.

## Author contributions

Conceptualization, SS, BYH, and SBMY; Data Collection, MZW, and BYH; Formal Analysis, SS and SBMY; Writing—Original Draft, SS and SBMY; Writing— Review and Editing, SS, BYH, and SBMY; Funding Acquisition, BYH, and SBMY.

## Data and Code availability

The datasets generated and/or analyzed during the current study are available at https://github.com/SangkyuSon/VRmaze. Correspondence and requests for materials should be addressed to SS and SBMY. The data could be shared upon the reasonable request to the corresponding authors.

## METHODS

### Subjects

Two male rhesus macaques (*Macaca mulatta*) were subjects in the current experiment. The University Committee on Animal Resources at the University of Rochester and the University of Minnesota approved all animal procedures. Animal procedures were designed and conducted in compliance with the Public Health Service’s *Guide for the Care and Use of Animals*.

### Surgical Procedures

A small prosthesis head fixation was used. Animals were habituated to laboratory conditions and then trained to perform oculomotor tasks for liquid rewards. We placed a Cilux recording chamber (Crist Instruments) over the area of interest (see *Recording Site* section for breakdown). We verified positioning by magnetic resonance imaging with the aid of a Brainsight system (Rogue Research). Animals received appropriate analgesics and antibiotics after all procedures. Throughout both behavioral and physiological recording sessions, we kept the chamber with regular antibiotic washes and we sealed them with sterile caps.

### Recording Site

Two Cilux recording chambers (Crist Instruments) were placed over areas 11 of OFC and 29 and 30 of RSC (Supplementary Figure 3.1C-D of reference; Öngür & Price, 2000) (**Figure 1B**). The position was verified by magnetic resonance imaging with the aid of a Brainsight system (Rogue Research Inc.) for subject P and a Cicerone system (developed by Dr. Matthew D. Johnson at the University of Minnesota) for subject S. Neuroimaging was performed at the Rochester Center for Brain Imaging, on a Siemens 3T MAGNETOM Trio Tim using 0.5 mm voxels. We confirmed recording locations by listening for characteristic white and gray matter sounds during recording, which matched the loci indicated by the Brainsight or Cicerone systems.

### Electrophysiological Techniques

Multicontact electrodes (V-probes, Plexon, Inc) were lowered until positioned within the OFC and RSC. Following a settling period, all active cells were recorded. Electrodes were lowered using a microdrive (NAN Instruments) until the target region was reached. This lowering depth was predetermined and calculated using either Brainsight or Cicerone to ensure that most of the V-probe contacts were in the recording region’s gray matter. Individual action potentials were isolated on a Ripple Grapevine system (Ripple, Inc.). Neurons were selected for study solely based on isolation quality; we did not pre-select based on task-related response properties. Cells were sorted offline manually by MZW with Plexon Offline Sorter (Plexon, Inc.). No automated sorting was used.

We confirmed the recording location before each recording session using our Brainsight system with structural magnetic resonance images taken before the experiment. Neuroimaging was performed at the Rochester Center for Brain Imaging on a Siemens 3T MAGNETOM Trio Tim using 0.5 mm voxels. We confirmed recording locations by listening for characteristic sounds of white and gray matter during recording, which in all cases matched the loci indicated by the Brainsight/Cicerone system with an error of ∼1 mm in the horizontal plane and ∼2 mm in the z-direction.

### Task

The task was designed by MZW and collaboratively coded and tested with the professional video game developer Mr. Benjamin Kalb (**Figure 1A**). A video of one subject performing the task can be found at https://www.haydenlab.com/maze. The task was controlled via a joystick with a speed ceiling. The joystick was custom-made for nonhuman primates. The joystick maintained complete directional motion freedom. No correction or assistance for joystick movement was provided to the monkeys. Joystick training was conducted first with a series of gradually more complex basic training programs designed by MZW to achieve joystick control proficiency. Then, monkeys were trained in the task virtual reality maze environment with standard shaping procedures. Before training on this task, subjects had been trained on other laboratory tasks in the lab, specifically, a sequence task (Blanchard & Hayden, 2014) and a standard economic choice task (Mehta et al., 2019; M. Z. Wang et al., 2022).

The maze was a virtual space, with both x- and y-coordinates ranging from -35 to 35 in arbitrary units. In the maze, participants were limited to navigating only within paths (width of 2-unit distance). Fences encircled the paths, preventing any attempts to cross over. Four paths connected the north and south part of the maze, and another four paths connected the east and west. The paths intersected vertically with each other, resembling the layout of Manhattan streets. Fourteen junctions (choice points) were present, with the center points of each junctions having x- and y-coordinates of (-21, -21), (-7, -21), (+7, -21), (+21, -21), (-21, -7), (-7, -7), (+7, -7), (+21, -7), (-21, +7), (-7, +7), (+7, +7), (+21, +7), (-7, +21), and (+7, +21), respectively. The overall arena of paths was similar to Tolman’s maze(Tolman & Honzik, 1930). As the subject navigates through the maze, they may encounter one of seven subgoals positioned at the coordinates (-14, -7), (0, +7), (-14, +21), (+7, +28), (-28, -21), (+14, -7), and (-28, +7). Subjects may reach the main goal, which was randomly located at the beginning of each session and remains until the end of the session. A small reward (0.125 ml) was provided upon reaching subgoals, and the *jackpot* reward (1.0 ml) was provided upon reaching the main goal. Rewards were provided when the subject reached a location in proximity to the main goal or subgoals within an invisible boundary (2-unit distance). The goals could not be seen from a far distance and were only visible when subjects were closer than a certain distance (7-unit distance). The goals were represented as a semi-transparent colored sphere placed at the designated location. After the subject was rewarded for reaching the main goal, the screen briefly blacked out, and then the subject was teleported to a random location. The teleportation location was set to be at least one junction-to-junction distance away (14-unit distance). Subjects P and S completed a total of 517 and 642 trials, encountering a total of 5194 and 3904 junctions (choice points) during 8 and 9 recorded sessions, respectively.

### Behavioral Features

The time window of the choice points (junctions) was defined when the subject’s position was within a radius of corridor width (2-unit distance) from the center of the choice point. From each choice point window, twelve low-level features of the subject’s behavior were extracted (**Figure 2C** and **Supplementary** Figure 3): (i) head turn speed (mean angular speed of directional change in the trajectory), (ii) head turn consistency (mean angular velocity of directional change in the trajectory), (iii) residence time at the choice point, spatial speed (iv) at the choice point, and (v) 500 ms before and (vi) after the choice point, joystick press strength (vii) at the choice point, (viii) 500 ms before and (ix) after the choice point, and spatial acceleration (x) at the choice point, (xi) 500 ms before and (xii) after the choice point.

Additionally, we extracted decision-related features at each choice point, including informativeness and optimality of the option paths. To compute the amount of information, at each junction (choice point), the number of previous visits to nearby junctions (i.e., potential choices) was respectively counted. Then, each visit count was divided by the total visit count of all potential choices. Shannon’s entropy was computed as the amount of *information* using the normalized visit count (i.e. *log*_2_(1/*p*), where *p* is the normalized number of previous visits). If the subject rarely or never chose a specific path, the information value would be high in that particular path. Among path options, we defined the path options with maximum information as an *informative* path and the non-maximum information path as a *not-informative* path.

Meanwhile, the optimality index measures how closely the subject approaches the reward if a particular path is taken, specifically defined as;

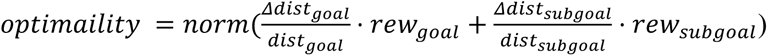

*rew_goal_* and *rew_subgoal_* indicate the amount of juice reward the subject gets if the subject reaches the main goal or the closest subgoal (i.e., 1 ml and 0.125 ml, respectively), and dist_goal_, dist_subgoal_, Δdist_goal_, and Δdist_subgoal_ denotes the Manhattan distance from the current location to main goal or the closest subgoal, and the changes in the Manhattan distance to the main goal or the closest subgoal if a certain path is chosen. The optimality index was normalized to a scale ranging from 0 to 100 (in percentage) so that the optimality index values over 50 indicate that the subject will be closer if choosing the path, and values below 50 indicate the subject will be at a farther location if choosing the path. Thus, we defined a particular path as a *not-optimal* path if the optimality was lower than 50, and an *optimal* path if higher than 50. Conceptually, the optimality index means the proportion of optimal choice if a certain path is chosen. The level of optimality and information among option paths was ranked (**Figure 2E**); first for the highest value in informativeness or optimality, second for the second-highest, and so forth, based on their respective values.

### Latent State Inference

With choice points in twelve low-level feature dimensions, we registered a k-means algorithm. Due to the different scales across twelve features, we normalized each feature dataset by dividing it by the standard deviation across the choice points. For each session, we trained a k-means clustering algorithm using choice points in 12-dimension from all the other sessions (training dataset) to estimate the learned centroids of the k-means clustering algorithm (leave-one-session-out procedure). Then, we measured the average distance of the corresponding session’s choice points (test dataset) and the closest trained centroid (within-cluster distance), or the second closest centroid (across-cluster distance). The across-cluster distance was subtracted from the within-cluster distance to estimate the model’s performance. Lower values indicate higher performance. The subtraction of across-cluster distance prevented the distance from merely decreasing by increasing the number of *k*. This procedure was repeated after resampling the choice points dataset from each session many times (bootstrapped), and after changing the number of *k* clusters from one to five (**Figure 2B**).

Then, we inferred the latent state in the sequential train of twelve behavioral features using the hidden Markov model (Rabiner, 1989). The number of hidden states was fixed to two, pre-identified by the k-means clustering algorithm. We assumed the two latent states would result from Gaussian mixtures of twelve low-level features. At each choice point, the hidden Markov model estimated a latent state and generated model-based behavioral outputs in a twelve-feature dimension. The model-based behaviors were fitted to the actual behavioral feature dataset using the Expectation-Maximization algorithm. The estimated hidden state for each choice point was used for further analysis. For the visualization purpose, we employed t-distributed stochastic neighbor embedding (t-SNE; van der Maaten, 2008) and isometric Mapping (Isomap; Tenenbaum et al., 2000) on the twelve-dimensional features (**Figure 2A** and **Supplementary** Figure 1)

### Linear-Nonlinear-Possion model (LNP)

The feature selectivity of RSC and OFC cells (i.e., tuning) was identified using the LNP model(Hardcastle et al., 2017). We assumed that each cell activity is the summation of filter-weighted features after an exponential nonlinear transformation (**Supplementary** Figure 4A).

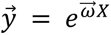

where 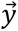, 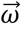, and *X* represents the row vector of predicted firing rates, the row vector of estimated filter weight, and the matrix of feature indices (respectively in size of 1 by *o*, 1 by *f*, and *f* by *o*; *o*, number of observations; *f*, number of feature measurements). We used 3 types of features: the subject’s spatial position, head direction, and spatial speed. The x- and y-coordinates of the subject were binned into 9 equal bins, respectively, for the spatial position, head directions into 4 (north, east, west, and south), and spatial speed into 7 (total 29 feature measurements). The feature indices matrix (*X*) was composed of one-hot column vectors for all observations. In each one-hot vector, the position, head direction, and speed of the corresponding observation were marked as one, and the values for the other features were set to zero. The filter weight (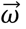) was fitted to match the Poisson process of the predicted firing rate (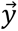) with the actual spike count. The firing rate changes as a function of feature measurements (i.e., tuning curve of place, head direction, and speed) were measured from both actual firing rate (true tuning) and model-based firing rate (model tuning).

To test if a particular cell was tuned to a specific feature measurement (a certain place, head direction, or speed), we measured the r-square value of the model tuning over the true tuning. The r-square values were then compared against the null distribution of the r-square (**Supplementary** Figure 4C). The null distribution was generated after randomly shuffling the relationship between a specific feature measurement and a firing rate (i.e., the columns in X were shuffled). If a cell’s r-square values were greater than the 99 percentile of the null distribution, the cell was counted as significantly tuned to the features (**Supplementary** Figure 4D). Cells were considered tuned to the spatial position feature only if they exhibited significant tuning to both x- and y-coordinate features.

### Goal Reactivation

We estimated the firing rate at each choice point and the firing rate within the time window when the subject reached the goal location in the lastest trial (i.e., the nearest rewarding experience). To prevent a single cell with high firing rates from dominating the result, the firing rates were normalized by subtracting the mean firing rate of choice points and then dividing by the standard deviation (z-scored). Then, Spearman correlation coefficients were measured between the normalized firing rate when reaching the goal location and at choice points as the amount of the goal reactivation (**Figure 3B**).

We measured the cell’s selectivity to informativeness (**Figure 3F**). For each cell, we measured a regression slope by using the information earned from choice points as the independent variable (x-axis) and the corresponding firing rate as the dependent variable (y-axis). We also repeated the same procedures after shuffling the order of the independent variable to earn a null distribution. If the regression slope fell outside the 95th percentile of the null distribution, we considered the cell to possess an information-selective property. Lastly, we considered a cell to possess goal-selective property if the mean firing rate when reaching the goal location fell outside the 95th percentile of the distribution of firing rate at other positions. To measure each cell’s role in goal reactivation, we employed an ablation approach; if excluding a specific cell led to a decrease in goal reactivation, then we can infer that the cell played a positive role in enhancing goal reactivation (**Figure 3G**). We counted the number of cells whose goal reactivation decreased independently after ablation for the cells with information-selective properties and cells with goal-selective properties.

### Path Reactivation

At each choice point, Spearman correlation coefficients were measured between the normalized firing rate at the current location and all the other possible choice points and corridors. Here, the corridor was defined as the time window between one choice point and the other choice point. To minimize the confounds of temporal drift, we only used datasets from choice points close to the current time (within 100 choice points). The coefficient values of the last occupied corridor and the choice point were averaged as *past reactivation* (**Figure 3E**). The coefficient values were rearranged into a 2D maze structure (reactivation map). Each value in the reactivation map represents the degree of reactivation for the corresponding location. The correlation coefficients of the current location were excluded. Subsequently, the reactivation maps were rotated to ensure that the vectors connecting the main goal location and the subject’s positions would point in the same direction. If the reactivation map was rotated at a slanted angle (not a multiple of 90 degrees), the nearest value was filled (nearest interpolation method). The rotated reactivation maps were aligned to the main goal location and averaged across to show *path reactivation* (**Figure 3C and D**).

The change in reactivation of the chosen path corridor was quantified by comparing the previous visit to the subsequent visit at the same location (**Figure 4C**). Before comparison, the average reactivation of non-chosen paths was subtracted from the reactivation of the chosen path for each visit. This subtraction ensures that the comparison between the previous visit and the subsequent visit was not driven by baseline differences. The percent change in the chosen path reactivation was quantified based on the absolute value of the previous reactivation.

### Agent-based model

For simplification, we first discretized the main task’s maze into a 9-by-9 grid miniature maze, where each bin denotes either a junction or corridor (see **Figure 3C** for an example). The subgoals were located in the same place as the actual task, and for each simulated session, the main goal was randomly located in one of the junctions. In each trial of a simulated session, an artificial agent was positioned on a junction that was a minimum distance away (= 2 junctions). The agent selected one of the option paths probabilistically in proportion to the weight. The path’s weight was decreased to 0.1 times its original value only if the agent never visited the path previously in the simulated session. Each choice point initially started with equal weight for all optional paths. For each visit to a specific choice point, the weight was updated with a pre-determined learning rule. The weight updates for one choice point were independent of those for the other. The weight of the *not-informative* path (see *Behavioral features* section for definition) was multiplied by (1-x) if the predetermined learning rule was x% devaluation for lack of information. The weight of the *informative* path was multiplied by (1+y) if the learning rule was y% reinforcement for information. The same weight update method was applied to the *optimal* and *non-optimal* paths. The weights of choice points were updated until the end of the simulated session, and reset when a new session began. Each trial simulation ended when the agent successfully reached the main goal location, and then the agent was teleported to a random location for the next trial. Each session consisted of 30 trials (i.e., 30 times of reaching the main goal), and we repeated 50 sessions. The agent’s performance (reward rate) was assessed as 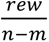, where *rew*, *n*, and *m* represent the acquired reward, the number of traveled junctions, and the minimum restarting distance from the main goal (i.e., 2), respectively. We iteratively assessed the reward rate of agents across gird combinations of learning rules on path informativeness and path optimality (**Figure 4D**).

### Inter-areal analysis

For each day, the inter-areal association between one area’s firing rate and the other area’s goal reactivation was measured using a generalized linear model approach as follows;

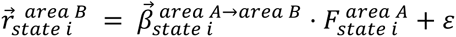

where 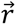, 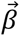, *F*, and *ε* denote the row vector of area B’s goal reactivation in either the explore-like state or exploit-like state, the row vector of the inter-areal association weight of neurons in area A, and the matrix of area A’s firing rate of the corresponding state, and scalar baseline term (respectively in size of 1 by *o*, 1 by *n*, *n* by *o*, 1 by 1; *o,* number of observation; *n*, number of neurons in area A). The 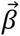s were independently measured for each state and each inter-areal relationship.

For each state, we measured the significance of 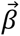s (**Figure 5C**). We first computed the cosine similarity between 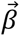 and 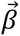*_bootstrapped_*, which was obtained after resampling the observation from all choice points, and then averaged them. Second, we computed the cosine similarity between 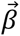 and 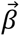*_random_*, which was obtained after randomly shuffling the relationships between 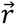 and *F* (i.e. shuffling the columns of *F*, then estimating 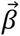*_null_*), to yield the null distribution. We compared the results of the first step with the null distribution of the second step.

The cosine similarity between 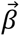*_explore-like_* and 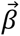*_exploit-like_* was also measured. The estimated similarity was compared against a null distribution of cosine similarities. The null distribution was generated with arbitrarily selected two 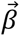s through the random resamplings of 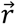s and *F*s (**Figure 5D**).

**Supplemental Video 1. Example of a subject performing the task, Related to Figure 1**. Subject’s view (top panel) and bird’s eye view (bottom panel). In the bird’s eye view, the location of the main goal, subgoal, and subject is indicated by a pink circle, a white circle with an ‘X’ inside, and a white circle with an arrow inside.

