## Supplementary Figures for "State-dependent Online Reactivations for Different Learning Strategies in Foraging"

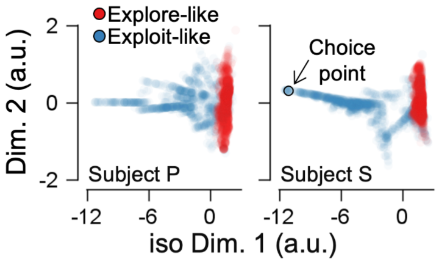


**Supplementary Figure 1. Visualization of low-level features in isomap space**. Each dot represents a value of a choice point. Color denotes the latent states in the hidden Markov model (blue: exploit-like; red: explore-like).

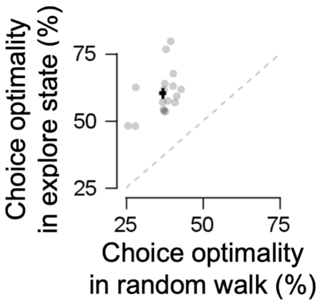


**Supplementary Figure 2. The optimality in the explore-like state was compared against the random walk**. The average optimality of option paths at each choice point was measured as the expected optimality in the random walk.

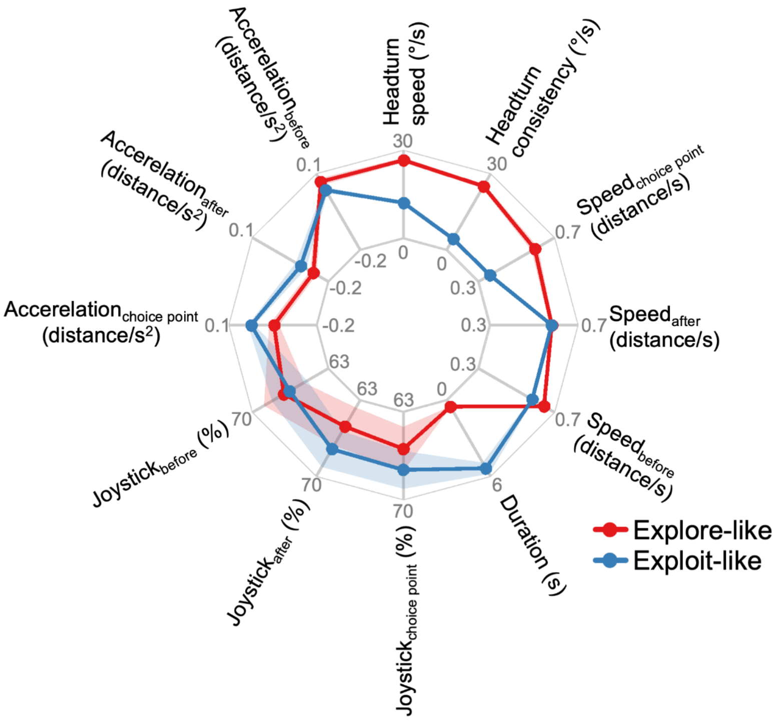


**Supplementary Figure 3. Twelve low-level features of two states**. Subscript words–choice point, after, and before–indicate the values during the subject at a choice point, 500 ms before and 500 ms after the choice point, respectively.

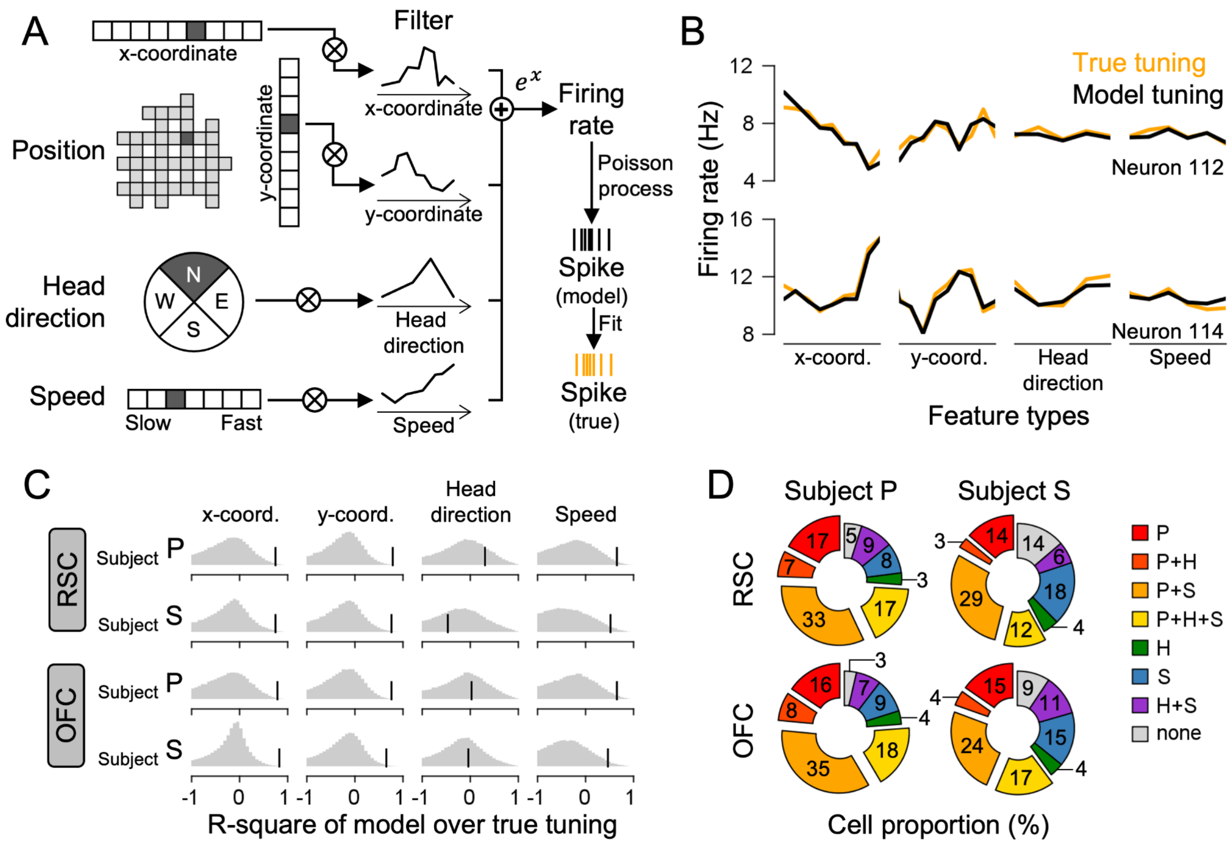


**Supplementary Figure 4. The result of the Linear-Nonlinear-Poisson (LNP) model analysis in RSC and OFC**. **(A)** A cartoon illustration of the LNP model. The x- and y-coordinate of the subject was discretized into 9 bins, head directions into 4 (north, east, west, and south), and speed into 7. The current value was marked as a filled bin in a one-hot vector. Then, the one-hot row vector was multiplied with the column vector of the learned filter, followed by exponential nonlinearity transformation. The Poisson process of the firing rate (model spike) was fitted to match the actual spike dataset. **(B)** Example cells' firing rate tuning curve (true tuning) overlaid with the LNP model's firing rate tuning (model tuning). **(C)** Average r-square values of the model tuning (black vertical lines) against the null distribution (gray background distributions). The null distribution was created by shuffling the matching relationship between one-hot vectors and their corresponding firing rates. **(D)** The proportion of significantly tuned cells (values inside indicate percentages). The cell was counted as tuned to a feature if the r-square value of the feature was greater than the 99 percentile of the null distribution. The spatial position tuning of a cell indicates that both x- and y-coordinate features were significant. P, H, and S represent the tuned features: position, head direction, and speed, respectively. The symbol '+' indicates that the cells were tuned to multiple features.

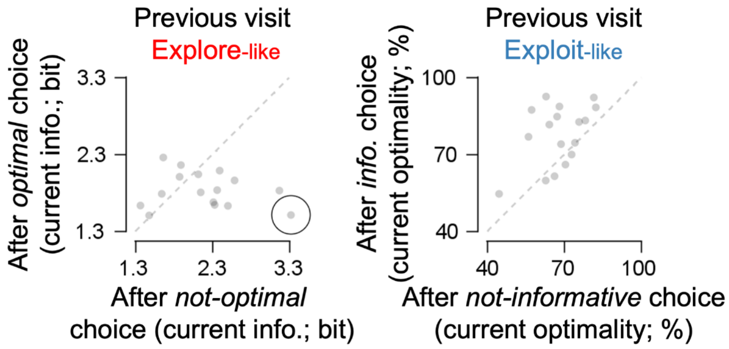


**Supplementary Figure 5. Subjects were not sensitive to results that did not align with their latent objective**. Following a choice made in an explore-like state, the information obtained during a subsequent visit remained consistent regardless of whether the preceding choice was optimal or suboptimal (subject P, $t_{(7)} = -1.555$, $p = 0.163$; subject S, $t_{(8)} = -1.664$, $p = 0.134$). Similarly, the subsequent choice exhibited comparable optimality, regardless of whether the previous choice in an exploit-like state was informative or uninformative (subject P, $t_{(7)} = 2.149$, $p = 0.068$; subject S, $t_{(8)} = 2.663$, $p = 0.028$). For visualization purposes, an extreme data point was plotted in a black circle (actual coordinate$_{(x,y)}$ = (8.4, 1.8)). These results suggest that subjects were not sensitive to their choice optimality in the explore-like state and choice informativeness in the exploit-like state.
